# The Germline Variants rs61757955 and rs34988193 are Predictive of Survival in Low Grade Glioma Patients

**DOI:** 10.1101/497354

**Authors:** Ajay Chatrath, Manjari Kiran, Pankaj Kumar, Aakrosh Ratan, Anindya Dutta

## Abstract

**Abstract:** Low grade gliomas are invasive brain tumors that are difficult to completely resect neurosurgically. They often recur following resection and progress, resulting in death. Although previous studies have shown that specific germline variants increase the risk of tumor formation, no previous study has screened many germline variants to identify variants predictive of survival in glioma patients. In this study, we present an approach to identify the small fraction of prognostic germline variants from the pool of over four million variants that we variant called in The Cancer Genome Atlas whole exome sequencing and RNA sequencing datasets. We identified two germline variants that are predictive of poor patient outcomes by Cox regression, controlling for eleven covariates. rs61757955 is a germline variant found in the 3’ UTR of *GRB2* associated with increased *KRAS* signaling, *CIC* mutations, and 1p/19q co-deletion. rs34988193 is a germline variant found in the tumor suppressor gene *ANKDD1a* that causes an amino acid change from lysine to glutamate. This variant was found to be predictive of poor prognosis in two independent low grade glioma datasets and is predicted to be within the top 0.06% of deleterious mutations across the human genome. The wild type residue is conserved in all 22 other species with a homologous protein.

*Implications:* This is the first study presenting an approach to screening many germline variants to identify variants predictive of survival and our application of this methodology revealed the germline variants rs61757955 and rs34988193 as being predictive of survival in low grade glioma patients.

## Introduction

Grade II and grade III (low grade) gliomas are primary brain tumors that are derived from glial cells and include astrocytomas and oligodendrogliomas. They are most commonly found in the cerebral hemispheres. They are highly invasive and therefore difficult to completely resect neurosurgically without significant patient morbidity. Following surgery, patients are typically treated with chemotherapy and radiation, though these tumors typically recur or progress to grade IV gliomas and are fatal.^1^ The median survival following low grade glioma diagnosis is around 7 years.^2^

While the 2007 World Health Organization’s (WHO) classification of central nervous system neoplasms differentiated between neoplasms primarily based on histological features, the updated 2016 WHO classification system now utilizes both molecular and histological parameters.^1^ Isocitrate dehydrogenase mutation (*IDH*) status, 1p/19q co-deletion status, telomerase reverse transcriptase (*TERT*) promoter mutation status, *MGMT* promoter methylation, *TP53* mutation status, and *ATRX* mutation status may be used to molecularly characterize gliomas.^1,3^ The availability of genomic data from patient glioma samples from groups such as The Cancer Genome Atlas (TCGA), the Chinese Glioma Genome Atlas (CGGA), and the Ivy Glioblastoma Atlas Project has substantially contributed to our understanding of these tumors.^4,5^

Many studies have utilized these datasets to identify gene expression signatures, microRNA expression patterns, somatic mutation status, and imaging characteristics that are predictive of survival in low grade gliomas.^6–8^ While studies have shown that germline mutations can increase an individual’s susceptibility for specific cancers,^9–12^ including a recent study that identified 853 pathogenic or likely pathogenic germline variants found in 8% of 10,389 cancer patients,^13^ no study has comprehensively screened all of the germline variants in a given cancer type to discover the prognostic variants in that cancer type. Although germline mutations have been shown to be prognostic in breast cancer^14^ and medulloblastoma^9^ in genes that have been well-characterized in the context of these cancers, these variants were not identified using an unbiased approach that screened a large number of germline variants. Identifying prognostic germline variants is challenging due to the limited effect size of germline variants, the large number of germline variants, and confounding clinical factors that may be associated with germline variants. Here we present a novel methodology for identifying prognostic germline variants and report two germline variants that we have found to be associated with survival in low grade glioma patients.

## Methods

### Glioma Datasets

491whole exome sequenced normal blood samples (WXS normal), 503 whole exome sequenced tumor samples (WXS tumor), and 501 RNA sequenced tumor samples (RNA tumor) from TCGA low grade glioma^4^ patients available on the Cancer Genomics Cloud (CGC)^15^ platform were used as part of this analysis. The clinical information was downloaded directly from the TCGA data portal using the GenomicDataCommons (https://bioconductor.org/packages/release/bioc/html/GenomicDataCommons.html) R package available through Bioconductor. Additional molecular characteristics about these TCGA patients were acquired by downloading the supplement from Ceccarelli et. al.^16^ The raw sequencing data from the Chinese Glioma Genome Atlas patients was downloaded using accession number SRP027383 from the Sequence Read Archive. Clinical information for these patients was downloaded directly from the project’s website (http://www.cgga.org.cn/).

### Variant Calling

Variant calling was performed on the TCGA low grade glioma whole exome sequenced normal blood samples (WXS normal), whole-exome sequenced tumor samples (WXS tumor), and RNA sequenced tumor samples (RNA tumor) using VarDict^17^ on CGC. The VarDict settings were set at default except for requiring mapping quality greater than 30, base quality greater than 25, a minimum of 3 variant reads, minimum allele frequency of 5%, and the removal of duplicate reads. We compiled a list of all of the unique variants and ran ‘samtools^18^ depth’ on all sequencing files requiring a mapping quality greater than 30. We determined the status of each variant in each patient from the three datasets (WXS normal sample, WXS tumor sample, and RNA tumor sample). The variant status at positions with fewer than ten reads for a given patient was changed to unknown. We used the WXS tumor samples to insert variant calls into the WXS normal samples at positions at which a variant status was listed as unknown in the WXS normal samples. If the variant status was still missing in a given patient, we then used the RNA tumor sample to insert variant calls into the combined WXS variant call set, allowing us to create the combined set of variant calls.

The same program parameters and approach were used to variant call and process the CGGA RNA sequencing dataset. All computation on the CGGA dataset was performed locally and not on CGC.

### Quality Control

We used annovar^19^ to determine the allele frequencies of the variants called by VarDict as listed in gnomAD (http://gnomad.broadinstitute.org/). We calculated the allele frequency of the variants in our study using the following formula:

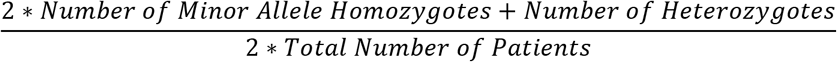

The R package GGally (https://cran.r-project.org/web/packages/GGally/index.html) was used to calculate the correlation between the four variant call sets and to display their correlations with each other. Only variants with an allele frequency of greater than 5% in gnomAD and found in 15 or more of the TCGA low grade glioma patients were tested for an association with survival by Cox regression.

Because we used the WXS tumor and RNA tumor samples to fill in missing variant calls, we evaluated whether somatic mutations were affecting the validity of our results. We first determined the percentage of variants called in the WXS tumor sample that were somatic mutations. To do this, we downloaded the set of somatic mutations generated by the TCGA Research Network.^20^ We then calculated the number of somatic mutations called in each patient in this variant call set and divided that number by the total number of variants called in that patient’s WXS normal sample. To assess whether somatic mutations were affecting the integrity of our results, we counted the number of times that a somatic mutation called by the TCGA Research Network overlapped with the set of germline variants that we were testing for an association with survival.

Since we used the RNA tumor sample to fill in missing variant calls, we evaluated whether RNA editing was having a significant impact on our analysis. To do this, we downloaded the set of over 2.5 million known RNA editing sites from a rigorously annotated database of RNA editing sites, RADAR.^21^ We counted the number of times that the germline variants that we were testing for an association with survival overlapped with any of the known 2.5 million RNA editing sites.

### Principal Component Analysis

In order to calculate principal components that could separate patients on the basis of race, we used PLINK^22^ to create a pruned set of germline variants to avoid bias from variants in linkage disequilibrium. Pruning was performed using a window size of 50 variants and a variance inflation factor of 2. These variants were used to calculate principal components using base R.

### Cox Regression and Receiver Operator Characteristic Curves

Lasso in the R package glmnet^23^ was run on 17 covariates (**Table 1**). Information about patient age, gender, tumor location, grade, treatment site, and TP53 mutation status was acquired from the TCGA data portal, while data for patient somatic mutation count, percent aneuploidy, TERT expression, IDH mutation status, 1p/19q co-deletion status, MGMT promoter methylation status, and chromosome 7 gain with chromosome 10 loss status was acquired from Ceccarelli et. al.^16^ The principal components were calculated as described above. 11 of these 17 covariates were selected for inclusion in the final model for survival prediction. The R packages survival^24^ and survminer^25^ were used to run Cox regression and create Kaplan-Meier curves. For each minor allele, we our model tested whether the minor allele was associated with a difference in survival outcomes with respect to the reference allele. False discovery rate correction was performed through Bonferroni correction.

**Table 1.** List of variables that are known to be associated with differences in survival in low grade glioma patients. 11 variables (red) were selected by Lasso for inclusion in the survival model. We used these 11 variables as covariates in our Cox regression model when testing each germline variant.

| Covariate |  | Median (Min-Max) or Number of Patients |
| --- | --- | --- |
| **Age** |  | 41 (14 - 87) |
| Gender | Female | 250 |
|  | Male | 200 |
| **Somatic Mutation Count** |  | 50 (0 - 12255) |
| **Percent Aneuploidy** |  | 11% (5.2E-4% - 95%) |
| log(TERT Expression) |  | 1.0 (0.0 - 9.1) FPKM |
| Principle Component 1 (PC1) |  | 0.043 (-0.091 - 0.064) |
| Principle Component 2 (PC2) |  | -0.017 (-0.23 - 0.17) |
| **Principle Component 3 (PC3)** |  | -0.53 (1.34E-4 - 0.33) |
| Histological Type | **Astrocytoma** | 172 |
|  | Oligoastrocytoma | 113 |
|  | Oligodendroglioma | 165 |
| Tumor Location | Frontal Lobe | 265 |
|  | Temporal Lobe | 125 |
|  | Parietal Lobe | 42 |
|  | Other | 18 |
| Grade | G2 | 212 |
|  | **G3** | 237 |
|  | Cannot Be Assessed | 1 |
| Treatment Site | **Henry Fords Hospital** | 82 |
|  | Case Western St. Joes | 90 |
|  | Other | 278 |
| IDH Mutant | Wild Type | 83 |
|  | **Mutant** | 367 |
| 1p/19q Co-deletion | Absent | 303 |
|  | **Present** | 147 |
| MGMT Promoter Methylation | Unmethylated | 80 |
|  | **Methylated** | 370 |
| Chr 7 gain/Chr 10 loss | Absent | 399 |
|  | **Present** | 51 |
| TP53 Mutant | Absent | 232 |
|  | Present | 218 |

Receiver operator characteristic (ROC) curves were created and evaluated using the survivalROC (https://cran.r-project.org/web/packages/survivalROC/survivalROC.pdf) and pROC (https://cran.r-project.org/web/packages/pROC/pROC.pdf) R packages. In order to test whether rs61757955 significantly improves the survival model consisting of the eleven covariates selected by Lasso, we compared the two ROC curves using the bootstrap method with 1000 iterations. We also used this bootstrapping approach to determine whether ANKDD1a expression levels, GRB2 expression levels, rs61757955, and rs34988193 together improve the survival model with respect to the eleven covariates selected by Lasso.

### RNA-Sequencing Data Processing

We downloaded the HTSeq FPKM quantification files for each patient from the Genomic Data Commons data portal. We only used gene quantification files from primary tumor samples as part of this analysis. Replicate samples from a single patient were averaged.

### Variant Correlation to Covariates and Somatic Mutations

In order to test for associations between the germline variants and genomic and histological tumor characteristics, we divided patients based on their germline variant status. We used the Wilcoxon rank-sum test to test for significant differences in each of the continuous variables between patients with and without a given variant. We used Fisher’s exact test to test for differences in each of the discrete variables using a similar approach. Somatic mutation calls were downloaded from Ellrott et. al.^20^

### Gene Set Enrichment Analysis

Gene set enrichment analysis (GSEA) of mRNA changes associated with rs61757955 and rs34988193 was performed by dividing the patients into two groups for each variant based on whether or not they had the reference allele at the position of the variant. For each germline variant, we calculated the log fold change for all genes expressed greater than one fragment per kilobase per million mapped reads (FPKM) between patients with the variant and without the variant. For each gene, fold change was calculated by dividing the median expression of the gene in patients with the variant by the median expression of the gene in patients without the variant. We used the log fold change to rank the genes from greatest log fold change to smallest log fold change. This file was used as input for GSEA.^26^

### Variant Annotation

In order to identify deleterious mutations, we annotated all variants by combined annotation dependent depletion (CADD) scores and only analyzed the variants predicted to be within the top 0.1% of all deleterious variants (CADD > 30).^27^ This led us to identify rs34988193 in *ANKDD1a* as a potentially deleterious variant predictive of survival. Because rs34988193 causes an amino acid change from positively charged lysine to negatively charged glutamate, we ran a BLASTp (httpps://blast.ncbi.nlm.nih.gov/Blast.cgi) search so that we could determine how many species have a protein homologous to ANKDD1a and how consistently the wild type lysine residue was conserved. We identified homologous sequences in 22 other species. These sequences were aligned using ClustalW in MEGA.^28^ We also annotated this variant with its PhyloP score.^29^ Because the crystal structure for *ANKDD1a* was not available, we downloaded the predicted model for this protein from Modbase (https://modbase.compbio.ucsf.edu/modbase-cgi/index.cgi) and calculated the Gribskov score using prophecy on EMBOSS.^30^ We retrieved linked variants from Ensembl using the population of Utah residents with Northern and Western European ancestry which is demographically similar to the TCGA low grade glioma patient population.

## Results

### Identification of High Quality Germline Variants

Our variant calling pipeline is shown in **Figure 1**. Briefly, we used the variant caller VarDict on Cancer Genomics Cloud to identify variants from whole exome sequencing (WXS) and RNA sequencing samples in about 500 low grade glioma patients. In total, we found 4,453,701 unique variants. We used ‘samtools depth’ to determine the sequencing depth at each of these variants for each patient and changed the variant status to ‘unknown’ for patients with sequencing coverage less than 10 reads at a given position. We created a set of combined variant calls by using the WXS and RNA tumor samples to fill in unknown values in the whole exome sequenced normal samples that resulted from having a sequencing coverage of less than 10 reads at a given position. This approach increased our sample size and enabled us to include many more variants in our analysis than if we had solely used variant calls from the whole exome sequenced normal blood samples. Ultimately, this left us with four sets of variants – WXS normal, WXS tumor, RNA tumor, and a combined set that resulted from merging the other three variant call sets, giving preference to the WXS normal and then WXS tumor variant calls. We used the combined variant call set when testing variants for an association with survival. We only tested variants found in 15 or more low grade glioma TCGA patients and listed in gnomAD as having an allele frequency of greater than 5%.

**Figure 1.**
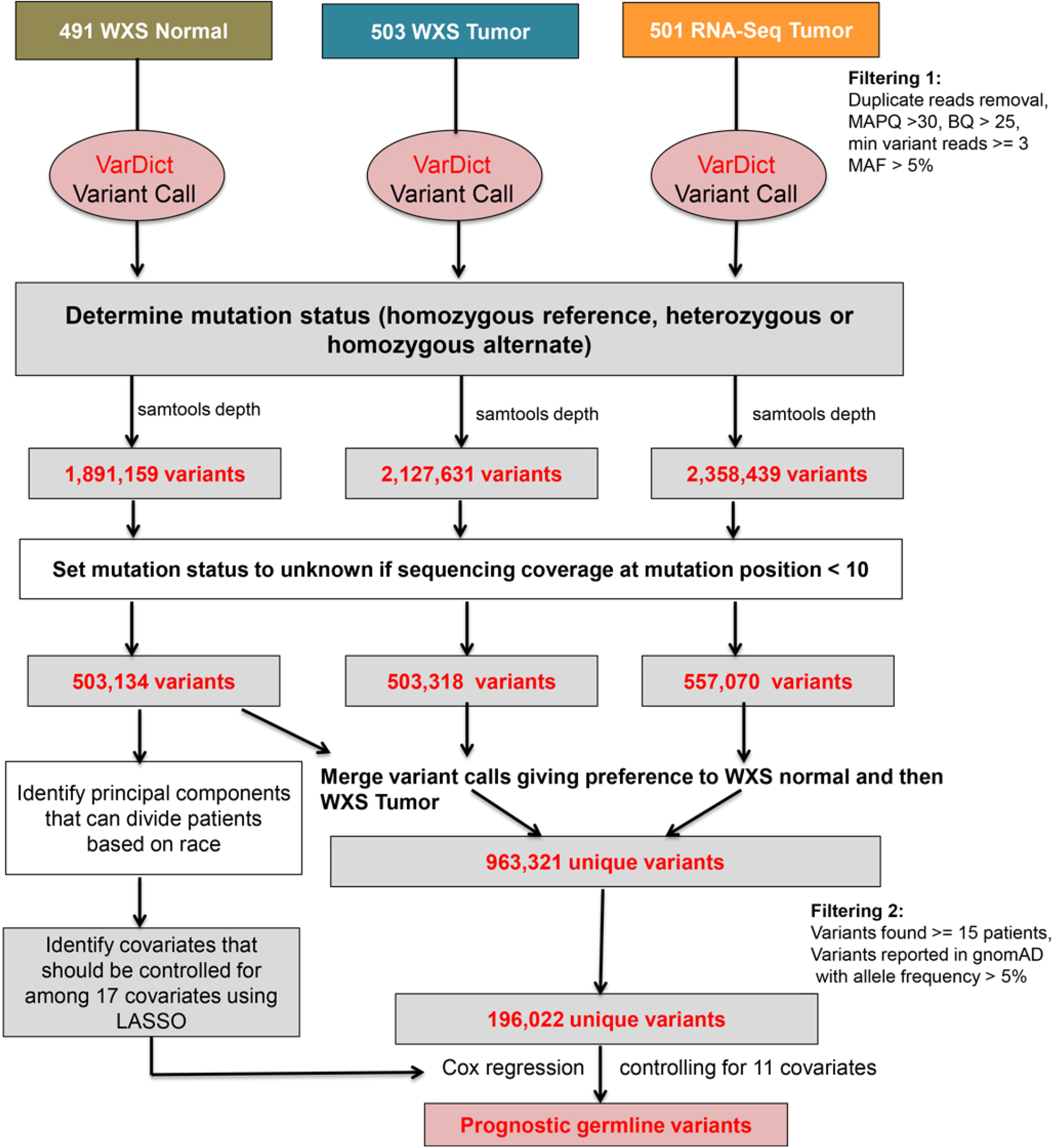
A flowchart describing the steps involved in identifying prognostic germline variants.

### Tumor Variant Calls are not Significantly Affected by Somatic Mutations or RNA Editing After Filtering

Because we used sequencing data from the WXS tumor and RNA tumor samples to fill in missing calls in the WXS normal samples, we evaluated our variant calls for contributions from somatic mutations and RNA editing. We first showed that the majority of variant calls in the tumor sample are germline variant calls. To do this, we counted the number of somatic mutations called by the TCGA Research Network’s analysis in each patient and divided that number by the number of variants that we called in the WXS normal sample.^20^ The median number of somatic mutations called per patient was 39. The median number of variants called in the WXS normal sample was 95,794. We therefore estimated that over 99.9% of variants called in the WXS tumor sample consisted of germline variants and that the percentage of somatic mutations in the WXS tumor sample across all patients was quite small (**Figure S1**). Because somatic mutations rarely occur at the same position, we suspected that the number of somatic mutations included in our study was extremely small since we limited our analysis to variants found in 15 or more of the low grade glioma patients and found in gnomAD with an allele frequency of greater than 5%. Indeed, only one of the 196,022 variants that we tested overlapped with a somatic mutation. This somatic mutation occurred in only a single patient (**Table S1**). Ultimately, we did not find any evidence to suggest that somatic mutations were impacting the quality of our analysis.

We next determined whether RNA editing was affecting our analysis by downloading the 2.5 million known RNA editing sites from the rigorously annotated RNA editing database, RADAR.^21^ Only 215 of the 196,022 variants that we tested were located at a position that overlapped with a known RNA editing site. We did not find any of these variants to be prognostic as part of our analysis. We therefore did not find any empirical evidence to suggest that somatic mutations or RNA editing impacted our findings (**Table S1**).

Finally, we established that our four variant call sets (WXS normal, WXS tumor, RNA tumor, and combined) were concordant with each other by calculating the allele frequency of each variant called in the four sets and demonstrating a very strong correlation between all pairs of variants (r > 0.98 for all pairs, **Figure S2**). To further evaluate the quality of our variants calls, we calculated the frequency of each allele and compared it to the frequency of these alleles as listed in gnomAD. Our alleles frequencies were well correlated with gnomAD (r > 0.93 for all four variant sets, **Table S2**). As expected, the distribution of allele frequencies is negatively skewed as the majority of the identified variants are rare (**Figure S2**). We used the variants from the WXS normal samples to determine the principal components. As expected, these principal components effectively separate patients on the basis of reported race (**Figure S3**).

### Identification of 271 Prognostic Germline Variants that are Independent of Clinical Covariates

In order to identify clinically relevant germline variants, we restricted our analysis to variants found in at least 15 patients in the TCGA dataset and found in gnomAD with an allele frequency of greater than five percent. This restricted our analysis to 196,022 testable variants (**Figure 2A**). In order to reduce the risk of identifying variants that are prognostic because they are confounded by other covariates known to be associated with survival, we used the machine learning algorithm Lasso to determine which of 17 covariates should be controlled for in our Cox regression model. Lasso regression was useful in the screening of these 17 covariates because it penalizes models based on the number of coefficients, allowing for the elimination of less predictive coefficients from the model. The algorithm selected 10 covariates known to be associated with differences in survival in low grade glioma (age, somatic mutation count, percent aneuploidy, histological subtype of astrocytoma, tumor grade, treatment site, *IDH* mutation status, 1p/19q co-deletion status, *MGMT* promoter methylation status, chromosome 7 gain/chromosome 10 loss status) along with the third principal component that we calculated (**Table 1**). Although the first two principal components are more effective in stratifying patients on the basis of race than the third principal component, the selection of the third principal component over the first two suggests that the third principal component contributes more information to the survival model than the first two principal components. This third principal component primarily separates African Americans from each other, suggesting that a subpopulation of African Americans experienced worse clinical outcomes in this dataset compared to other groups. We ran Cox regression on all 196,022 variants one at a time, controlling for these 11 covariates, to identify germline variants predictive of survival.

**Figure 2.**
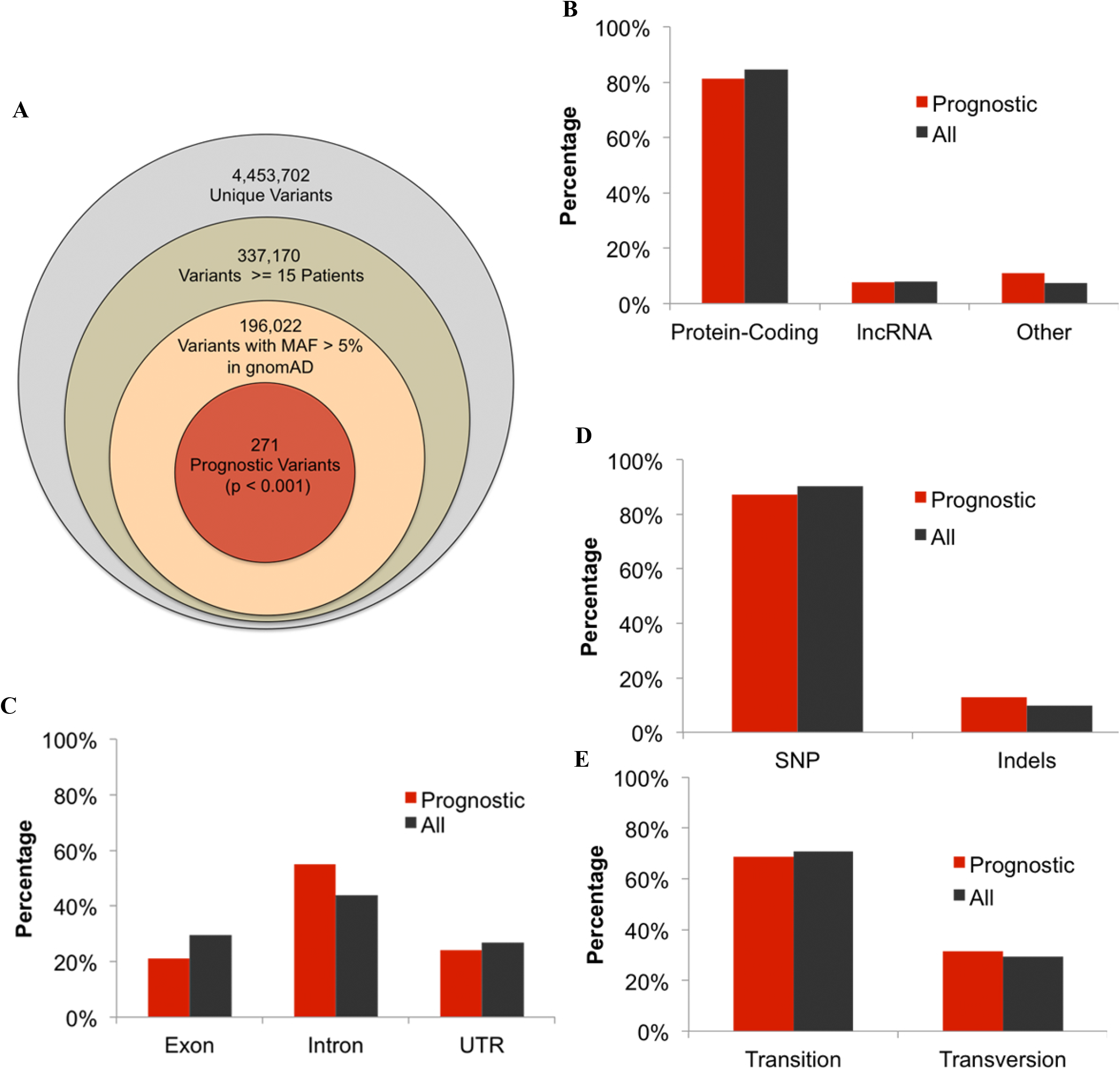
Prognostic germline variants in the TCGA dataset. **(A)** Of the 4.4 million unique variants called in the TCGA dataset, we ran Cox regression on the 196,022 germline variants found in gnomAD with an allele frequency greater than 5% and found in 15 or more of the TCGA low grade glioma patients. **(B-E)** Similar to the 196,022 germline variants, the 271 prognostic variants are mostly found in **(B)** protein-coding genes, **(C)** are located in introns, and are **(D)** single nucleotide polymorphisms (SNP). **(E)** Most single nucleotide polymorphisms cause transitions.

We identified 271 germline variants that are predictive of survival (p < 0.001) (**Figure 2A**). As is the case with germline variants in general, the majority of these germline variants are found in protein-coding genes (**Figure 2B**), are located in introns (**Figure 2C**), and are single nucleotide polymorphisms (**Figure 2D**). Most single nucleotide polymorphisms are transitions (**Figure 2E**).

### The Germline Variant rs61757955 in *GRB2* is Associated with Poor Prognosis

We identified two germline variants that are highly predictive of survival after false discovery rate correction (FDR < 0.10) (**Figure 3A**, **Table 2A**). rs61757955 results in a mutation in the 3’ UTR of Growth Factor Receptor Bound Protein 2 (*GRB2*) and is associated with a poor prognosis (p=7.08E-10, hazard ratio(HR)=20.4, **Figure 3B**, **Table 2A**). To determine whether rs61757955 enhances the survival model compared to the eleven clinical covariates alone, we calculated a risk score for each patient using a Cox regression model with rs61757955 and the other 11 covariates and a risk score using the 11 covariates alone. Using these risk scores, we determined the rate at which a patient would be correctly labeled as alive or dead at 7 years with a given false positive rate to create a receiver operator characteristic curve. The increased area under the curve suggests that rs61757955 enhances the survival model compared to the eleven clinical covariates alone (p=0.0489, **Figure 3C**). The allele frequency of rs61757955 is close to 0% according to the 1000 Genomes Project^31^ in the Chinese population and, as expected, did not show up in the Chinese Glioma Genome Atlas. We also found rs28672782, a germline variant found in the intron of BRSK2, to be associated with a favorable prognosis, though the testable sample size for this variant was small and the maximum follow up for patients with this variant was only three years. Therefore, we did not investigate this variant further (**Figure S4**, **Table 2A**).

**Figure 3.**
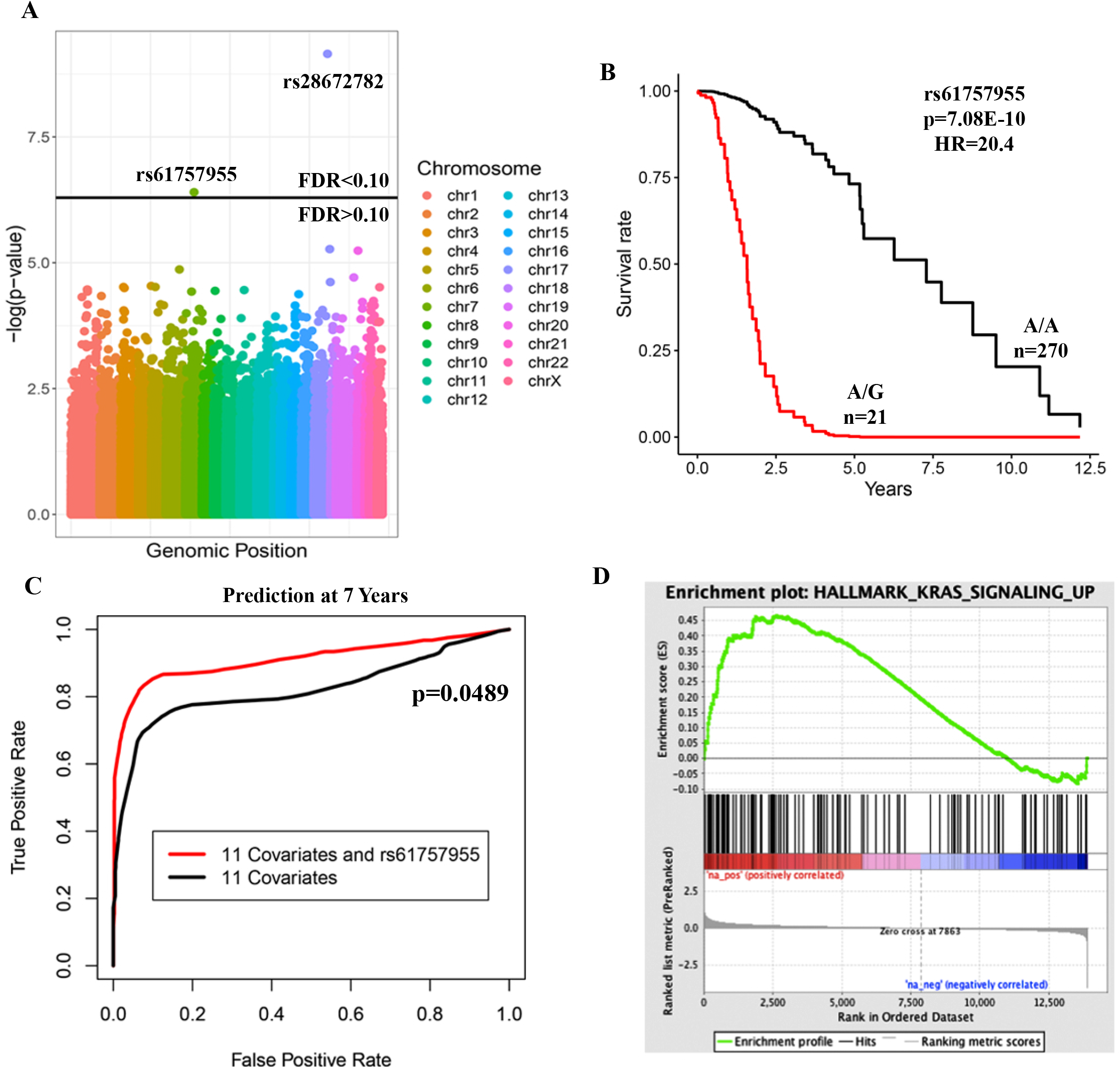
rs61757955 is a highly prognostic germline variant identified in the TCGA dataset. **(A)** Manhattan plot showing the p-values resulting from testing each germline variant by Cox regression, controlling for the 11 variables in red in Table 1. Two variants passed the FDR threshold in the TCGA dataset. **(B)** A Kaplan-Meier plot depicting the deleterious outcome associated with rs61757955, adjusting for the eleven covariates. **(C)** Receiver operator characteristic curve at 7 years. rs61757955 increases the area under the curve compared to the 11 covariates alone, suggesting that it improves the clinical model. **(D)** Separation of patients on the basis of whether or not they have this germline variant to determine which genes are induced (red) or repressed (blue) in patients with rs61757955. Subsequent gene set enrichment analysis reveals that patients with this germline variant exhibit upregulation of the genes involved with *KRAS* signaling.

**Table 2.** Description of the prognostic germline variants identified in this study. **(A)** A description of the two prognostic germline variants (FDR<0.10) in the TCGA dataset identified when testing all 196,022 germline variants. **(B)** A description of the prognostic germline variant (FDR<0.10) rs34988193 in *ANKDD1a* identified when the analysis was restricted to only germline variants with a combined annotation dependent depletion (CADD) score greater than 30 in the TCGA and CGGA datasets.

| Variant | Chrom | Pos | RefAlt | Population<br>Frequency | Sample<br>Size | Number of<br>Heterozygotes | Number of<br>Homozygotes | Gene Name | Median<br>Expression<br>(FPKM) | p-value | Hazard<br>Ratio |
| --- | --- | --- | --- | --- | --- | --- | --- | --- | --- | --- | --- |
| rs61757955 | 17 | 75318086 | A G | 5.01% | 291 | 21 | 0 | GRB2 | 42.2 | 7.08E-10 | 20.4 |
| rs28672782 | 11 | 1446622 | C T | 16.27% | 50 | 15 | 5 | BRSK2 | 7.29 | < 1E-16 | 1.15E-10 |

B
| Variant | Chrom | Pos | RefAlt | Population<br>Frequency | CADD<br>Score | PhyloP<br>Score | Gene<br>Name | Dataset | Sample<br>Size | Number of<br>Heterozygotes | Number of<br>Homozygotes | p-value | Hazard<br>Ratio |
| --- | --- | --- | --- | --- | --- | --- | --- | --- | --- | --- | --- | --- | --- |
| rs34988193 | 15 | 64943580 | A G | 30.90% | 32 | 8.42 | ANKDD1a | TCGA | 450 | 199 | 52 | 0.00113 | 1.73 |
|  |  |  |  |  |  |  |  | CGGA | 76 | 18 | 2 | 0.0743 | 1.79 |

In order to test whether rs61757955 in *GRB2* is associated with an increased risk of other genomic abnormalities, we separated patients on the basis of this variant to see if there was a difference in the incidence of the genomic or histological variables (**Table 3**). We found this variant to be associated with an increased incidence of 1p/19q co-deletions (p=0.038). Because 1p/19q co-deletions are frequently seen in Capicua transcriptional repressor (*CIC*) mutated gliomas^32^ and *CIC* aberrations are known to be a driver in low grade glioma tumorigenesis,^33^ we tested whether there was a difference in the incidence of *CIC* mutations in patients with this variant. 38% of patients with this variant had *CIC* mutated gliomas, whereas only 16% of patients without the variant had a *CIC* mutation (p=0.0168, **Table 3**). Although the incidence of oligodendrogliomas was elevated in patients with the variant compared to patients without the variant, consistent with reports from the literature that 1p/19q co-deletions and *CIC* mutations are enriched in oligodendrogliomas,^32^ this difference was not statistically significant (p=0.475). Since rs61757955 is in a non-coding region, we also tested whether this variant is associated with differences in gene expression. We separated patients based on their variant status and calculated the log fold change of each gene between patients with the variant and patients without the variant. This data was used as the input for gene set enrichment analysis (GSEA). We found rs61757955 to be associated with increased *KRAS* signaling (FDR=0.015) (**Figure 3D**).

**Table 3.** The association between the germline variant rs61757955 and genomic and histological variables. Patients were divided based on whether or not they had the germline variant rs61757955. Patients with the germline variant rs61757955 were more likely to have *CIC* mutated gliomas and the 1p/19q co-deletion.

| Variable | Mean or Percentage<br>(Wild Type) | Mean or<br>Percentage<br>(Mutant) | p-value |
| --- | --- | --- | --- |
| **CIC Mutated** | **15.9%** | **38.1%** | **0.017** |
| **1p/19q Co-deletion** | **25.2%** | **47.6%** | **0.038** |
| Oligodendroglioma | 33.7% | 42.9% | 0.475 |
| Total Somatic<br>Mutation Count | 30.9 | 30.0 | 0.766 |
| Percent Aneuploidy | 15.1% | 11.7% | 0.524 |
| Astrocytoma | 38.1% | 42.9% | 0.651 |
| Grade 3 | 53.0% | 42.9% | 0.497 |
| IDH Mutated | 78.1% | 85.7% | 0.583 |
| 1p/19q Co-deletion | 25.2% | 47.6% | 0.038 |
| MGMT Promoter<br>Methylation | 77.8% | 81.0% | 1.000 |
| Chr 7 Gain/Chr 10<br>Loss | 13.0% | 9.5% | 1.000 |
| Expression of<br>GRB2 (FPKM) | 45.7 | 44.4 | 0.636 |

Because we only have whole exome sequencing and RNA sequencing data from The Cancer Genome Atlas, we do not know whether the upregulation of genes in the *KRAS* signaling pathway and the increased incidence of *CIC* mutations and 1p/19q deletions are due to this variant or a linked variant in a regulatory region that we would be able to analyze with whole genome sequencing data. Therefore, we identified the four other variants that are genetically linked to rs61757955 in the European population, the population which is most similar to the TCGA low grade glioma patient population (**Table S3**). These variants did not pass the criteria to be included within the 196,022 testable variants that we had identified at the beginning of this study but could become useful in the future.

### rs34988193 is a Deleterious Germline Variant Present in *ANKDD1a* Associated with Poor Outcomes

In order to identify prognostic variants that are predicted to be deleterious due to effects on the encoded protein, we repeated our analysis but restricted it to only variants with a combined annotation dependent depletion (CADD) score greater than 30 and expression greater than one FPKM on average. 81 variants met this criteria. These variants correspond to the top 0.1% of deleterious mutations as predicted by this scoring system. We found the germline variant rs34988193 in the tumor suppressor gene *ANKDD1a* to be associated with poor prognosis in the TCGA dataset (p=0.001, HR=1.73, FDR < 0.10, **Figure 4A-B**, **Table 2B**). Because this variant is found in both the European and Asian populations, we were able to test whether this variant is also predictive of survival in the independent Chinese Glioma Genome Atlas (CGGA) dataset. We found this variant to be predictive of survival in the CGGA dataset and we found the hazard ratio that we calculated in CGGA to be very similar to the hazard ratio calculated in the TCGA dataset (p=0.0743, HR=1.79, **Figure 4C**, **Table 2B**). rs34988193 is not linked with any other variant in the European population. We did not find any enriched pathways after performing gene set enrichment analysis and this variant was not associated with differences of any of the genomic or histological variables (**Table S4**).

**Figure 4.**
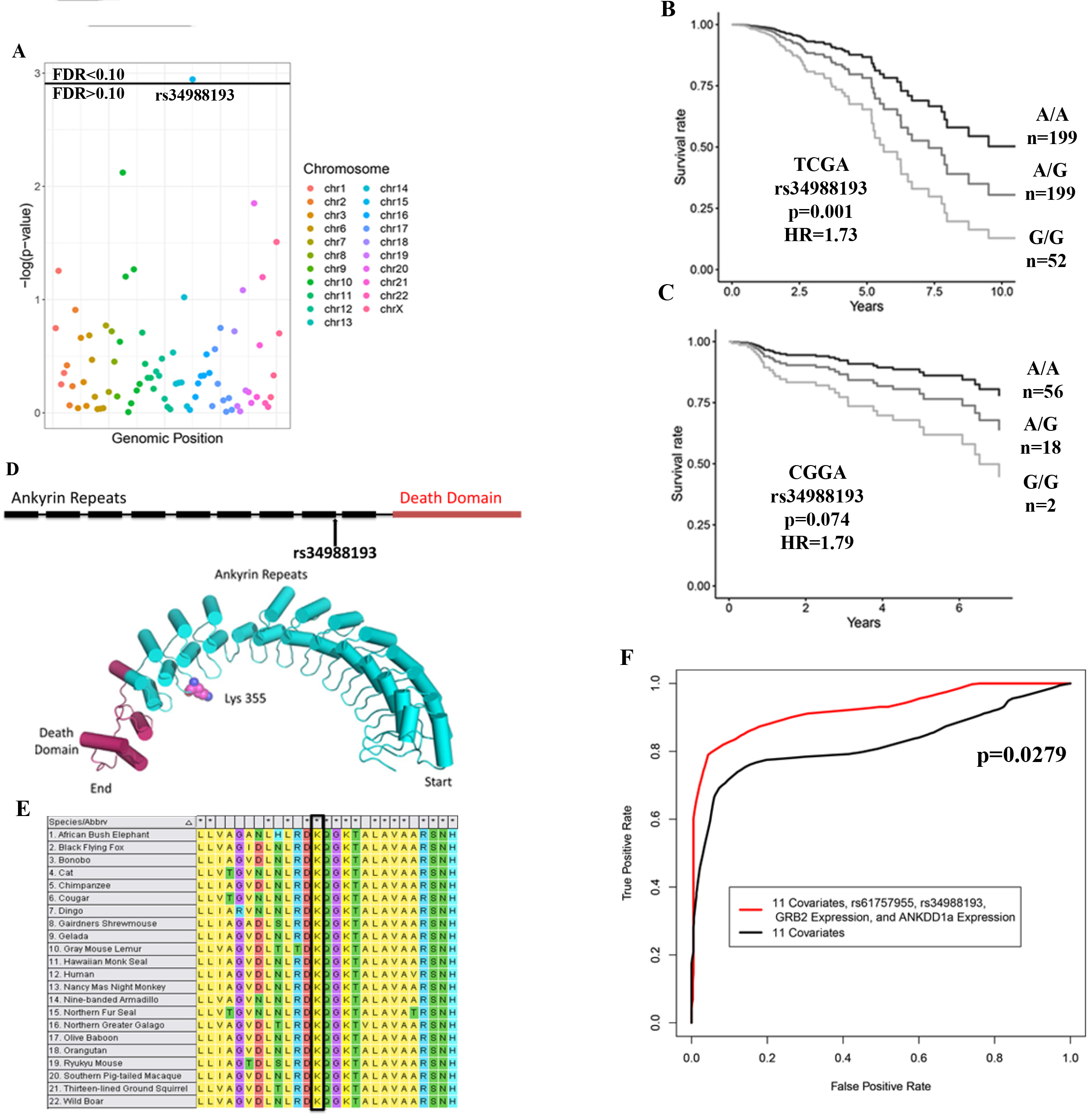
rs34988193 is a prognostic variant predicted to be highly deleterious. **(A)** A Manhattan plot with the p-values resulting from testing each germline variant by Cox regression, controlling for the eleven covariates in red in Table 1. rs34988193 is prognostic (FDR<0.10) in the TCGA when restricting the analysis to the top 0.1% most deleterious variants by combined annotation dependent depletion (CADD). **(B-C)** Kaplan-Meier plots depicting the deleterious outcome associated with rs34988193 in the **(C)** TCGA and **(D)** CGGA datasets, adjusted for the eleven covariates. **(D)** A schematic showing that this variant is located in the ninth ankyrin repeat of *ANKDD1a*. The predicted protein structure of *ANKDD1a* reveals that this variant leads to an amino acid change from lysine to glutamate on the loop of an ankyrin repeat. **(E)** Multiple sequence alignment of *ANKDD1a* in 22 species showing that lysine is conserved at this position in all of the species with this protein. **(F)** Receiver operator characteristic curves comparing the ability of two survival models to label patients as alive or dead after seven years of follow up. The inclusion of rs61757955 variant status, rs34988193 variant status, GRB2 expression and ANKDD1a expression significantly improves the survival prediction compared to the eleven covariates bolded in red from table 1 alone (p=0.0279).

*ANKDD1a* contains ten ankyrin repeat domains and one death-like domain. This variant causes a non-synonymous mutation in the last codon of the ninth ankyrin repeat domain. The AAG to GAG codon change results in the incorporation of negatively charged glutamate instead of the wild type positively charged lysine residue in the loop between ankyrin repeats nine and ten (**Figure 4D**). This variant has a CADD score of 32 and is therefore predicted to be in the top 0.06% of deleterious mutations across the human genome. We performed a BLASTp search using the *ANKDD1a* protein sequence to identify homologous sequences in 22 other species. We aligned these sequences using ClustalW and found that this lysine residue is conserved in all 22 of these species (**Figure 4E**). The PhyloP score at this position is 8.42, suggesting that evolution is occurring much more slowly than expected at this residue assuming no selection pressure. We determined the position-specific profile Gribskov’s score for a lysine to glutamate amino acid change at this position using the multiple sequencing alignment from 23 species to be 15 to 3, suggesting that this variant is highly unfavorable.

### Combined Model Predicts Survival Better Than Clinical Covariates Alone

As a result of this analysis, we found the germline variants rs61757955 in the 3’ UTR of GRB2 and rs34988193 in the protein-coding region of ANKDD1a to be predictive of survival in low grade glioma patients. We constructed a survival model consisting of the eleven clinical covariates, rs61757955, rs34988193, GRB2 expression, and ANKDD1a expression and generated a receiver operator characteristic curve by using this model to categorize patients as alive or dead after seven years of follow up. This combined model is significantly better at predicting survival compared to the eleven clinical covariates alone (p=0.0279, **Figure 4F**).

## Discussion

Up until this point, the identification of prognostic features in gliomas has been limited to clinical factors, somatic mutations, gene expression changes, and methylation pattern changes.^6–8^ Although many studies have commented on how germline variants could enable physicians to better individualize patient care by being able to better predict how a patient might respond to chemotherapeutic treatment,^34–36^ most large-scale studies have focused on identifying germline variants that predispose or protect an individual to a disease.^13,37^ These studies have not focused on understanding how germline variants can be used to individualize patient care following diagnosis. Identifying prognostic germline variants is difficult due to the large number of germline variants, the limited effect of any single germline variant, and clinical factors that may confound the effect of germline variants. In this study, we have developed a novel method that can be used to identify prognostic germline variants and we have used that method to identify two variants that are predictive of survival in the TCGA dataset. The germline variant rs61757955 in *GRB2* is not found in the Asian population and so could not be confirmed in an independent dataset. In contrast, the germline variant rs34988193 in *ANKDD1a* is found in both the European and Asian populations, and remarkably, was found to be prognostic with very similar hazard ratios in both the TCGA and CGGA datasets.

Studies of germline variants using TCGA datasets typically solely utilize the WXS normal blood samples.^13,38^ One major disadvantage to this approach is that it limits the analysis to genes within the capture regions of the whole exome sequencing kits used by the study.^4^ In this study, we combined the information from both the whole exome sequencing and RNA sequencing datasets for a given patient to identify germline variants outside of the whole exome sequencing capture region. Our approach had the added benefit of providing us with more information for a given variant for variants with low sequencing depth in the whole exome sequencing datasets. We do not believe that this approach significantly affected the accuracy of our variant calls because the allele frequencies calculated from the RNA sequencing dataset were well correlated with the allele frequencies from gnomAD and with the allele frequencies calculated from the whole exome sequencing datasets. We showed that somatic mutations and RNA editing did not affect the integrity of our finding. Only one somatic mutation in a single patient overlapped with the 196,022 variants that we tested in our analysis and only 215 of the 196,022 variants that we tested overlapped with the 2.5 million known RNA editing sites. We did not find any of these variants to be predictive of survival. Instead, we feel that the increased sample size resulting from the additional sequencing coverage greatly outweighs any effect that somatic mutations or RNA editing had on our results.

We next needed to devise an approach to using these germline variants in a Cox regression model. We first had to decide how to deal with the absence of a variant in the variant call file. The variant could be absent because the patient was wild type for that allele or because the sequencing depth at that position was too low to make the variant call. We therefore determined the sequencing depth of each variant at each position so that we could exclude patients with low sequencing depths for the testing of specific variants. Testing a large number of variants increased the probability of a variant being significant solely because it was confounded with another significant variable. To avoid this issue, we tested each variant while controlling for 11 other covariates that we found to be predictive of survival. In this study, we found rs61757955 to be associated with differences in 1p/19q co-deletion status. By including the 1p/19q co-deletion as a covariate in our model, we were able to estimate the effect of rs61757955 independent from the 1p/19q co-deletion status and the other ten covariates.

*GRB2* is a signal transduction adaptor protein that plays an oncogenic role in a variety of cancers.^39–42^ *GRB2* plays an important role in the *RAS/RAF/ERK* pathway. Its SH2 domain binds the phosphotyrosine of activated growth factor receptor, while its two SH3 domains bind the guanine nucleotide exchange factor son of sevenless (*SOS*) protein, resulting in *SOS* recruitment to the plasma membrane and subsequent *RAS* activation. *RAS* binds and activates the kinase *RAF*, which phosphorylates the kinase *MEK. MEK* phosphorylates and activates extracellular signal-regulated kinase (*ERK*) which transmits the signal to transcription factors in the nucleus. This results in cell proliferation.^43^ We found the variant rs61757955 located in the 3’ UTR of *GRB2* to be associated with poor prognosis in glioma patients. Separating patients on the basis of this variant revealed that the *KRAS* signaling pathway is upregulated in patients with this variant. As described above, *GRB2* plays a well-characterized role in this pathway.^43^ We also found this variant to be associated with an increased incidence of *CIC* mutations and 1p/19q co-deletions. *CIC* is a known driver of low grade glioma pathogenesis.^33^ Mutations in *CIC* are common in oligodendrogliomas and are associated with poor prognosis.^4,32^ Although patients with rs61757955 variant exhibited an elevation in the incidence of oligodendrogliomas which we expected given the increased incidence of *CIC* mutations and 1p/19q co-deletions,^32^ this difference was not statistically significant. It is possible that this germline variant or the four other germline variants that it is linked with increase a patient’s risk for oligodendrogliomas with the *CIC* mutation and 1p/19q co-deletion.

In this study, we were only able to study variants in the whole exome or RNA sequencing data. Although it is possible that the 3’ UTR of *GRB2* has regulatory activity or affects *GRB2* protein translation efficiency, it is also possible that one of the variants that rs61757955 is linked to regulates the *KRAS* signaling pathway. None of the four linked variants are in the protein coding sequence of *GRB2* so that if they upregulate RAS activity, like the rs61757955, they likely do so by regulating the expression of *GRB2*. While recent large-scale sequencing studies have published patient whole genome sequences,^44^ this data is not yet available for gliomas. We will be able to apply our approach to variants in regulatory regions in the future to specifically identify these prognostic variants when whole genome sequencing data for gliomas is available. Our inability to further study this variant in the CGGA dataset due to this variant being rare in Asian populations is a limitation of this study which could be addressed in the future with the availability of additional glioma sequencing datasets. This result also suggests that the clinical usefulness of specific germline variants is dependent on the frequency of that germline variant in the population.

*ANKDD1a* is a tumor suppressor gene that has been shown to inhibit cell autophagy and induce apoptosis in glioblastoma multiforme (GBM). It directly interacts with and upregulates *FIH1*, resulting in inhibition of *HIF1α* activity and decreased *HIF1α* half-life. This induces apoptosis in GBM cell lines in hypoxic microenvironments. Hypermethylation of this gene is common in GBM and leads to decreased *ANKDD1a* expression and increased cell proliferation.^45^ We found the germline variant rs34988193, located at the end of the ninth of ten ankyrin repeat domains in this protein, to be associated with a poor prognosis in low grade glioma patients in both the TCGA and CGGA datasets. The hazard ratio independently calculated using the two datasets is remarkably similar. The wild type lysine residue is conserved in all 22 species with a homologue to *ANKDD1a* and this position has a high PhyloP score. This variant is predicted to be within the top 0.06% of deleterious mutations in the human genome by CADD score^27^ because it causes a change from a positively charged lysine residue to a negatively charged glutamic acid residue in the loop of this ankyrin repeat. Ankyrin repeats are common domains known for their involvement with protein-protein interactions.^46,47^ Previous studies have suggested that mutations in the loops of ankyrin repeats may disrupt protein-protein interactions.^48–50^ The change from a positively to negatively charged amino acid resulting from the germline variant rs34988193 in the loop of *ANKDD1a* may disrupt *ANKDD1a*’s protein interaction partners and could explain the poor prognosis associated with this variant seen in two independent datasets. Given the amino acid change, further studies involving rs34988193 in *ANKDD1a* could be directed towards experimentally determining whether or not this variant alters *ANKDD1a*’s protein-protein interactions.

rs61757955 in *GRB2* and rs34988193 in *ANKDD1a* could also be used to enhance predictions made by survival models clinically, as we found that these variants are significant predictors of prognosis even after controlling for eleven covariates. The prognostic effect of rs34988193 in *ANKDD1a* seems to be fairly reliable, as we found that this variant had a similar hazard ratio in both the TCGA and CGGA datasets. Our approach could be used in the future to identify sets of germline variants that together enhance the predictions made by survival models, though the current number of low grade glioma sequencing samples is small relative to the large number of possible combinations of germline variants. Focused studies on particular sets of genes or pathways could potentially get around this low sample size problem by drastically limiting the number of variants studied. We believe that this study provides researchers with an effective approach to identifying biologically significant germline variants and provides clinicians with germline variants that could enhance currently existing survival models.

## Supporting information

## Acknowledgements

We thank Dr. Wei Min Chen at the University of Virginia for his statistical input on our Cox regression models. We thank Dr. Ana Damljanovic and Dr. Liz Williams for their assistance with computation on the Cancer Genomics Cloud Platform. We thank dbGAP for providing us with access to The Cancer Genome Atlas data. We thank all of the patients and their families that participated in The Cancer Genome Atlas and Chinese Glioma Genome Atlas studies. We thank the Dutta lab members for the valuable feedback during the drafting of this manuscript.

