## Supplementary material for "The Germline Variants rs61757955 and rs34988193 are Predictive of Survival in Low Grade Glioma Patients"

Figure S1

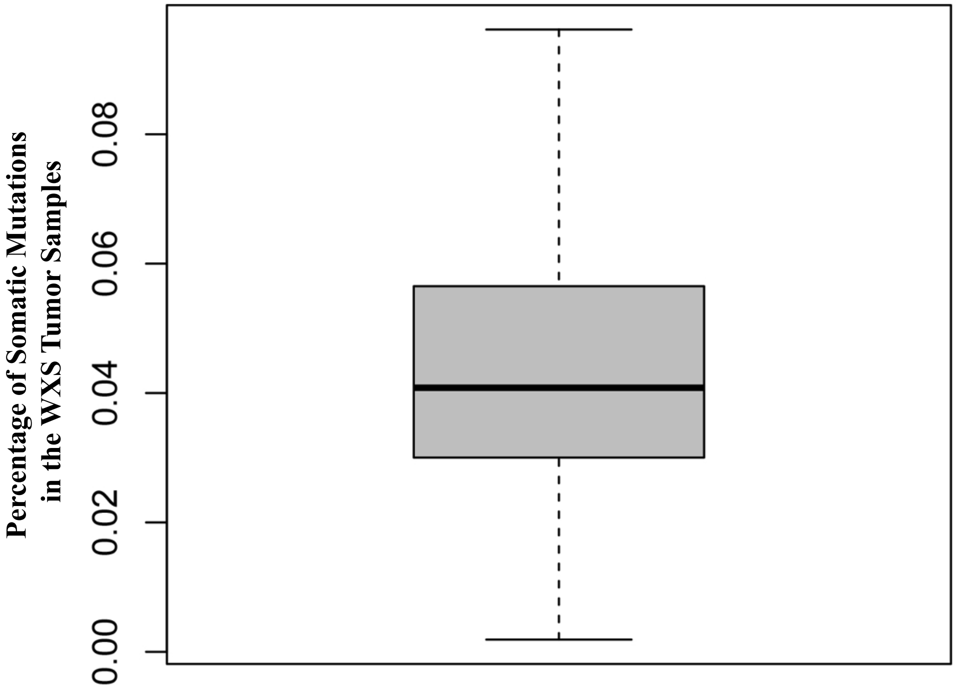


**Figure S1**. A boxplot representing the percentage of variants called in the whole exome sequenced (WXS) tumor sample that is likely somatic mutations. This value was calculated by counting the number of somatic mutations called in each patient by The Cancer Genome Atlas (TCGA) Research Network and dividing that number by the number of variants called in the WXS normal sample. Even before filtering, most variants called in the WXS tumor sample are germline variants.

**Figure S2**

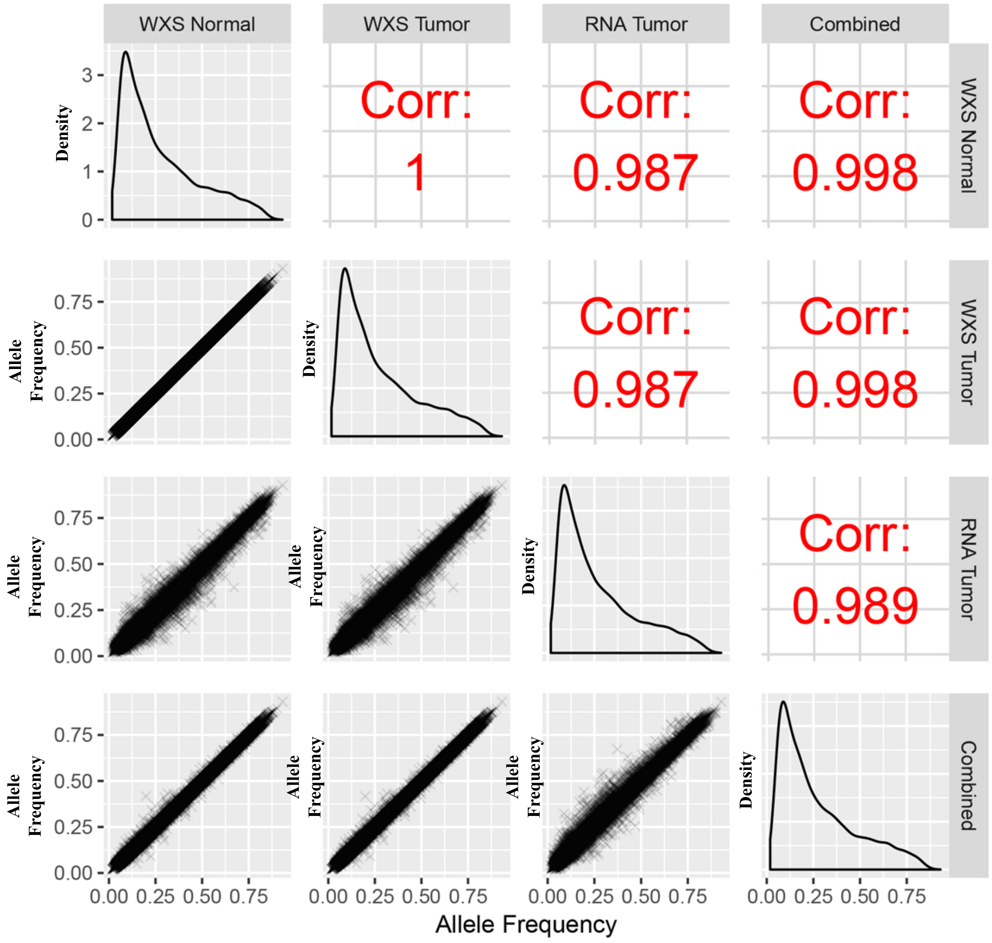


**Figure S2.** Correlation between the variant allele frequencies calculated from the four variant sets and the distribution of allele frequencies. The Pearson correlation panels on the top right (red) indicate that the calculated allele frequencies of the four variant sets are well-correlated with each other. This is depicted graphically in the bottom left panels with scatterplots. The distribution of allele frequencies is plotted along the diagonal. As has been shown in other studies, the distribution is negatively skewed because most minor alleles in the human population are rare.

**Figure S3**

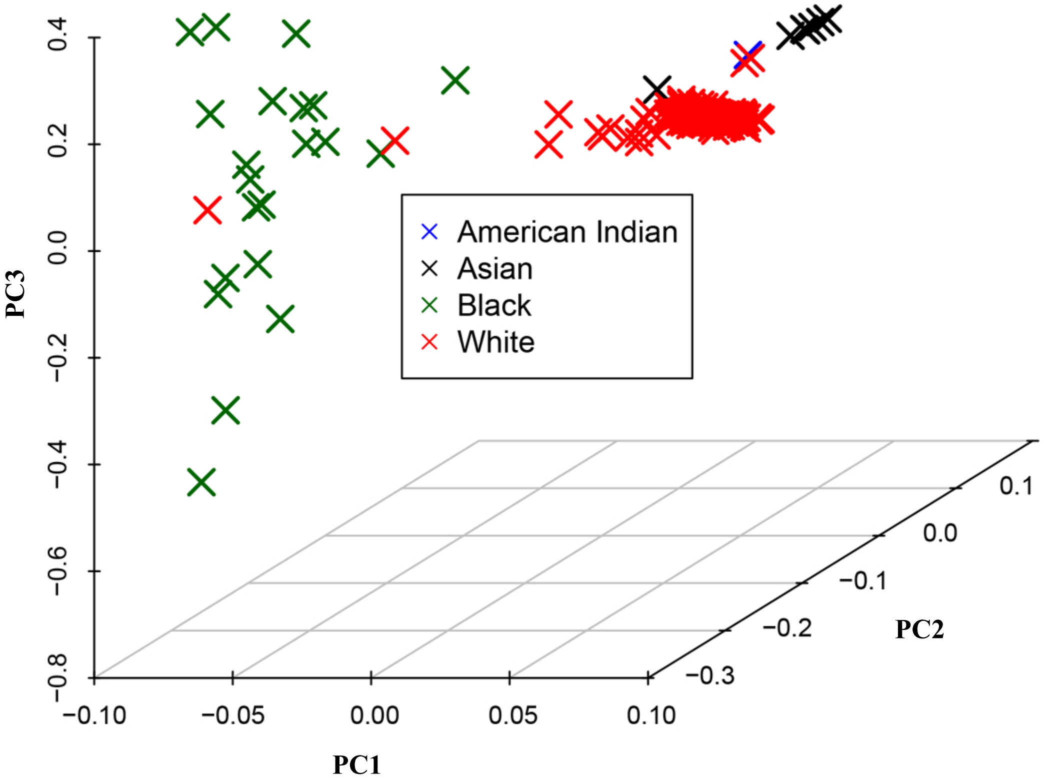


**Figure S3.** Principal components calculated from germline variants from the whole exome sequencing data from the non-tumor samples. Our principal components effectively stratify patients on the basis of patient-reported race.

**Figure S4**

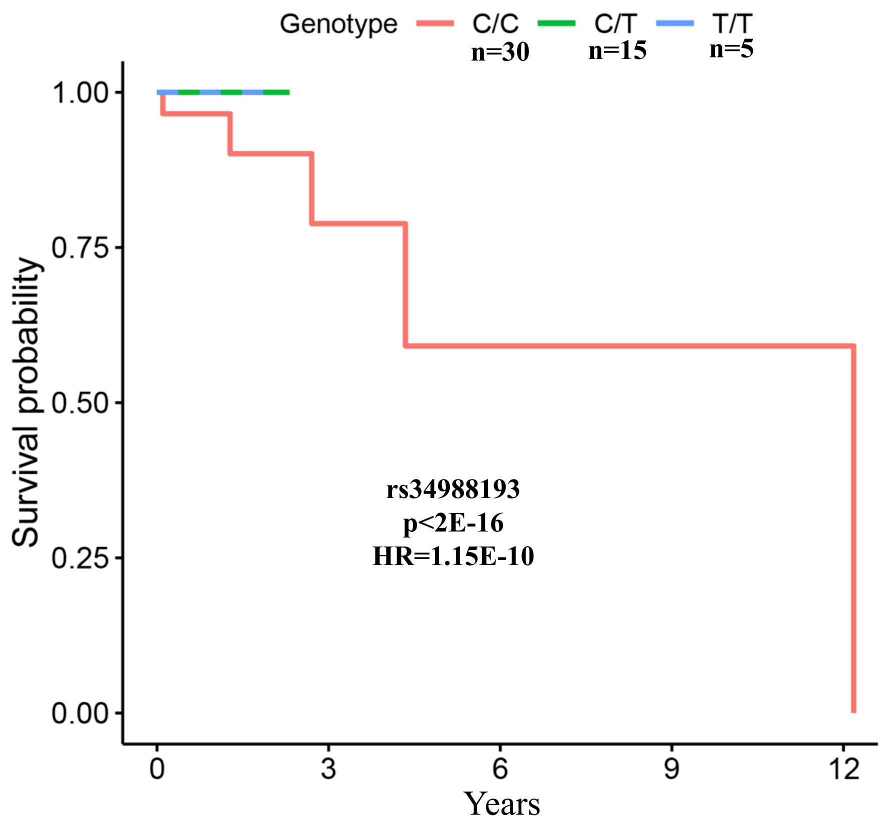


**Figure S4**. Kaplan-Meier plot for the germline variant rs28672782 in BRSK2. Although we found this germline variant to be prognostic (FDR<0.10), we decided not to further investigate this germline variant due to only 50 patients having sufficient sequencing depth at this position and due to the maximum follow up of patients with this germline variant being only three years.

Table S1

| Quality Check | Percentage |
|---|---|
| Percentage of Somatic Mutations Called in the WXS Tumor Sample | < 0.1% |
| Percentage of Somatic Mutations Included in this Analysis After Filtering | < 0.001% |
| Percentage of Germline Variants Included in this Analysis that Overlap with a Known RNA Editing Site | 0.1% |

**Table S1**. Quality control checks reveal that somatic mutations and RNA editing did not affect the results of our analysis. Less than 0.1% of variants called in the whole exome sequenced (WXS) tumor sample are somatic mutations in the TCGA low grade glioma patients. After our filtering steps, only one somatic mutation of the 196,022 variants tested in a single patient persists as part of our analysis. Of the 196,022 variants that we tested, only 0.1% of these variants overlap with a known RNA editing site. We did not find any of these variants as prognostic in our analysis, implying that somatic mutations and RNA editing did not compromise the quality of our analysis.

**Table S2**

| Variant Set | Pearson Correlation Coefficient with gnomAD |
|---|---|
| WXS Normal | 0.963 |
| WXS Tumor | 0.964 |
| RNA Tumor | 0.937 |
| Combined | 0.946 |

**Table S2**. Correlation between the allele frequencies calculated in our four variant sets and the allele frequencies reported by gnomAD. Our calculated allele frequencies are well-correlated with the allele frequencies reported by gnomAD, suggesting that our variant calls are high quality.

**Table S3. Variants genetically linked to rs61757955 in the European population**

| Variant | Chromosome | Position | Distance from rs61757955 (base pairs) | Correlation ($r^2$) | Location |
|---|---|---|---|---|---|
| rs56298430 | 17 | 75341627 | 23541 | 1 | Intron of GRB2 |
| rs41282071 | 17 | 75320799 | 2713 | 0.936 | Intron of GRB2 |
| rs55771008 | 17 | 75342476 | 24390 | 0.936 | Intron of GRB2 |
| rs72850335 | 17 | 75298213 | 19873 | 0.879 | 18,863 Base Pairs Upstream of GRB2 |

**Table S3.** Variants genetically linked to rs61757955 in the European population, the population that is most similar to the TCGA low grade glioma patient population. The upregulation of KRAS signaling may be due to rs61757955 or to one of these four linked variants.

Table S4

| Variable | Mean or Percentage (Wild Type) | Mean or Percentage (Mutant) | p-value |
|---|---|---|---|
| Total Somatic Mutation Count | 31.5 | 76.9 | 0.436 |
| Percent Aneuploidy | 13.4% | 15.1% | 0.466 |
| Astrocytoma | 41.7% | 35.5% | 0.204 |
| Grade 3 | 53.8% | 51.8% | 0.704 |
| IDH Mutated | 82.9% | 80.5% | 0.542 |
| 1p/19q Co-deletion | 28.6% | 35.9% | 0.107 |
| MGMT Promoter Methylation | 82.4% | 82.1% | 1.000 |
| Chr 7 Gain/Chr 10 Loss | 10.6% | 12.0% | 0.657 |
| Expression of ANKDD1a | 2.53 | 2.6 | 0.473 |

**Table S4.** Results from testing for an association between the germline variant rs34988193 and genomic and histological variables. We did not find the germline variant rs34988193 to be associated with any changes in genomic or histological variables by separating patients on the basis of whether or not they had the germline variant rs34988193.
